# Protein expression is amplified by stabilizing mRNA using gene-specific antisense oligonucleotides

**DOI:** 10.64898/2026.09.14.749961

**Authors:** Benjamin D. Boros, Aishwarya Nambiar, Miwei Hu, Dylan A. Galloway, Colin P. Florian, Courtney F. Jungers, Joseph D. Dougherty, Kathleen M. Schoch, Sergej Djuranovic, Timothy M. Miller

## Abstract

Effective and safe strategies to restore gene expression in haploinsufficiency disorders or to augment protective protein signaling have long been attractive but challenging therapeutic goals. Here, we demonstrate the potential of antisense oligonucleotides (ASOs) to upregulate gene targets through mRNA stabilization. The 3’ untranslated regions (UTRs) of mRNAs regulate stability via activating and repressive elements. One prominent destabilizing cis-element found in ∼30% of transcripts is the Pumilio RNA-binding protein recognition element (PRE), which can be targeted to enable gene-specific protein upregulation. A scan for PREs highlights hundreds of disease-relevant transcripts that may be targetable via ASO-mediated gene-specific blocking of Pumilio binding. We show that masking specific PREs with ASOs produces sustained protein upregulation through mRNA stabilization using multiple reporter and endogenous gene targets. Using PRE-masking ASOs, we neutralized PRE-mediated destabilization of TANK-binding kinase-1 (*TBK1*) mRNA and restored TBK1 protein expression in participant-derived fibroblasts with TBK1 haploinsufficiency, causal for amyotrophic lateral sclerosis. Additionally, by targeting PRE sites in the myoprotective gene follistatin (*FST*), we achieved effective *FST* mRNA upregulation in tested mouse tissues. These findings suggest that ASO strategies that neutralize destabilizing 3’UTR sequences like PREs may provide a powerful approach to upregulate haploinsufficiency-related genes or enhance protective protein levels for therapeutic benefit.

**One Sentence Summary:** Targeting inhibitory mRNA sequences with antisense oligonucleotides allows for controlled increase in gene and protein levels.

## INTRODUCTION

Inherited or acquired protein insufficiency underlies thousands of human conditions. Genetic haploinsufficiency, in which loss of a single gene copy is sufficient to cause disease, may contribute to >1,000 human disorders ^1^. For example, loss-of-function mutations in TANK-binding kinase 1 (*TBK1*) result in amyotrophic lateral sclerosis (ALS) and/or frontotemporal dementia (FTD) through gene haploinsufficiency ^2^. In addition, haploinsufficiency of *PKD1*/*PKD2* accounts for ∼85% of autosomal dominant polycystic kidney disease (ADPKD), which affects approximately 500,000 individuals in the United States ^3^. In these cases, as disease is caused by having 50% of the normal gene level, even a modest upregulation of protein towards normal levels may have substantial therapeutic impact. In addition to genetic haploinsufficiency, acquired insufficiencies in protein or protein signaling during aging, such as reduced muscle strength, underlie dozens of other conditions ^4,5^. Strategies that supplement lost DNA or mRNA with viral transfer approaches are promising but are challenged by difficulties with drug delivery, titratability, and risks of immune response ^6,7^. Thus, there remains a significant need for strategies to restore or augment gene expression and protein levels. We hypothesize that gene upregulation achievable by targeting mRNA regulatory regions could provide a safe, well-developed, and gene-specific therapeutic strategy.

The processing of nascent RNA from transcription to translation is an extensively regulated process that accounts for considerable variability in protein expression ^8,9^. Regulation of protein synthesis is influenced by activating and inhibitory controls on mRNA stability and translation efficiency, often coordinated by interactions of untranslated regions (UTRs) with RNA-binding proteins (RBPs) and small regulatory RNAs (e.g., microRNAs). The 3’UTR is the preferential binding site for many repressive RBPs that include Argonaute and Pumilio family proteins PUM1 and PUM2, with each conferring different degrees of inhibition. Argonaute is directed through short non-coding microRNAs to specific 3’UTR motifs, where this complex may repress target mRNA translation or stability on the order of ∼20% inhibition, but this varies widely across gene targets. PUMs are well-characterized repressors of mRNA stability and, to a lesser degree, translation that recognize a highly stereotyped 8 base pair 3’UTR motif, termed the Pumilio recognition element (PRE). About 40% of gene 3’UTRs contain at least one PRE, and the presence of a single PRE might repress protein synthesis by 30-50% depending on the gene target ^10^. This repression scales linearly with PRE burden, and ∼15% of genes contain two or more PREs. PUM proteins are ubiquitously expressed and repress PRE-containing mRNA targets by accelerating their decay, restricting cap-dependent translation, facilitating repression at nearby microRNA sites, or remodeling of the local secondary structure. Thus, we hypothesized that neutralizing PRE-mediated destabilization of individual genes might provide gene-specific protein upregulation.

We reasoned that gene upregulation achievable by targeting mRNA regulatory regions, such as with antisense oligonucleotides (ASOs), could provide a safe, well-developed, and gene-specific therapeutic strategy. ASOs are a nucleic acid-based technology that has achieved clinical success in treating monogenic illnesses. Examples include nusinersen for spinal muscular atrophy, tofersen for ALS linked to mutations in *SOD1*, inotersen for transthyretin amyloidosis, and mipomersen for familial hypercholesterolemia ^11,12^. ASOs are short (∼15–25 base pair) DNA- or RNA-like sequences that bind to target RNAs via sequence complementarity. Depending on their design, ASOs can modulate splicing (e.g., nusinersen) or degrade target mRNA by recruiting RNase H (e.g., tofersen) ^13^. More recently, ASOs have been explored for their potential to upregulate gene expression. Current ASO upregulation strategies involve degrading negative trans-acting regulators, disrupting inhibitory elements on target mRNAs, or redirecting non-productive alternative splicing products ^14–19^. While these approaches cover many genomic targets, they are often not broadly applicable or are difficult to identify and validate. In contrast, PREs, which are common, strongly repressive, and highly stereotyped, are amenable to neutralization by ASOs for gene-specific upregulation, primarily through mRNA stabilization.

Here, we validate PRE-masking ASOs and their application to multiple PRE-containing genes implicated in haploinsufficiency disorders. Importantly, we demonstrate the therapeutic potential of PRE-masking ASOs to haploinsufficiency disorders by amplifying expression of the ALS-related gene *TBK1* through *TBK1* mRNA stabilization. We apply this concept to other PRE-regulated genes associated with haploinsufficiency, *CDKN1B* and *DEPDC5*. Finally, we extend this strategy *in vivo* by targeting conserved PREs on the myoregenerative and protective gene *FST* (follistatin) ^20^, achieving effective upregulation in human cells and mouse liver. Collectively, PRE-masking ASOs introduce a novel antisense strategy to amplify gene expression with broad applications for haploinsufficiency-related disorders and protective signaling pathways.

## RESULTS

### Pumilio proteins regulate many genes associated with haploinsufficiency related diseases

Pumilio proteins are well-characterized repressors of mRNA stability, suggesting Pumilio - bound sequences as targets for gene upregulation. Therefore, we leveraged existing datasets to identify disease genes potentially regulated by Pumilio proteins (PUMs). In the subset of genes expressed in HEK293 cells, 3369 transcripts are bound by PUMs as determined by RNA-immunoprecipitation sequencing. The 2520 transcripts with overlap of “bound genes” and PRE-containing genes represents targets with high likelihood for regulation by PUM **(Figure 1A** and **Table S1)** ^10^. This set of genes contains ∼193 haploinsufficiency-associated genes (annotated by OMIM) and ∼682 other disease-associated genes (annotated by Orphanet). We selected six gene transcripts that are disease associated (*ITPR1*, *TBK1, DEPDC5, SLC2A1, LDLR* and *FST*) as potential targets ^2,21–25^. These targets are also strongly bound by Pumilio proteins PUM1 and PUM2 as determined by previously published RNA-immunoprecipitation sequencing results in HEK293 cells (**Figure 1B,C** and **Table S2**) ^26^. To test whether PUM proteins negatively regulate expression of these selected transcripts, we depleted PUM1 and PUM2 using siRNA in HEK293 cells. Indeed, PUM depletion increased mRNA of each gene target by >1.5-fold (**Figure 1D**).

**Figure 1.**
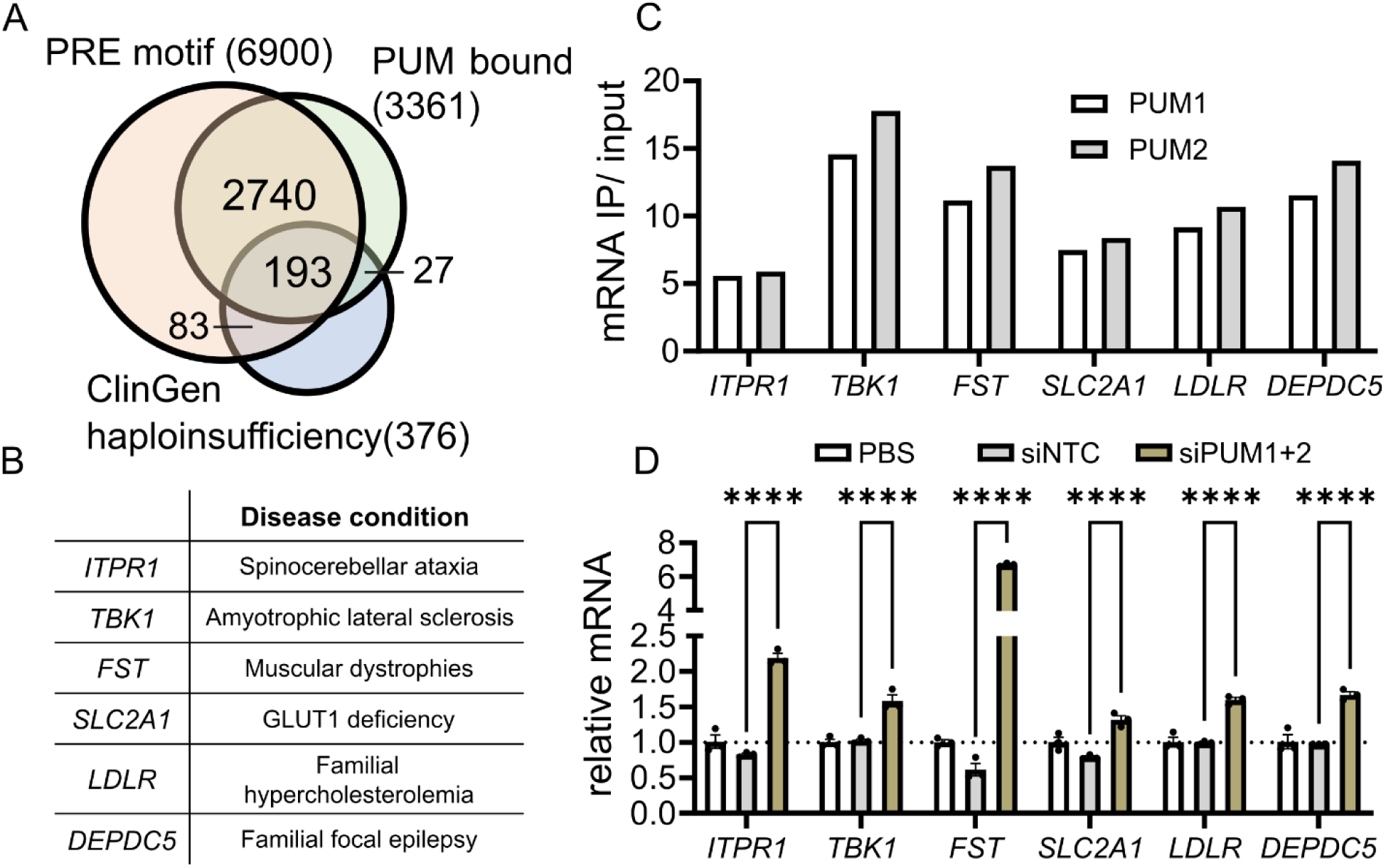
Pumilio proteins regulate thousands of transcripts associated with gene haploinsufficiency or protection against disease. **(A)** Venn diagram depicting the intersection of genes containing Pumilio regulatory element (PRE) motifs, mRNA that are bound by PUM proteins, and haploinsufficiency-associated genes. **(B)** Selection of gene targets associated with disease that contain PREs and are bound by PUM proteins. **(C)** Enrichment of selected mRNA targets with PUM proteins by RNA-immunoprecipitation sequencing (reanalysis of results from Yamada et al., 2017). **(D)** PUM proteins were depleted by siRNA for 48 hours in HEK293 cells, then mRNA was collected for RT-qPCR analysis of indicated targets. The mRNA target was normalized to the geometric mean of *GAPDH* and *ACTB*. (n=3 biological replicates per gene; Two-way ANOVA with Dunnett’s post-hoc test: all: p<0.001) ****p<0.0001.

Therefore, PUMs are putative targets for increasing protein levels, but targeting PUMs themselves would be predicted to upregulate many genes simultaneously.

### Masking PREs on 3’UTRs upregulates disease-associated protein targets

PUMs recognize and bind at a highly stereotyped 8 base pair motif (*UGUAHAWW*) that is readily identifiable within a gene 3’UTR (**Figure 2A**). We tested whether neutralizing Pumilio binding to a single gene target using antisense oligonucleotides (ASOs) could provide gene-specific upregulation by augmenting mRNA stability (**Figure 2B**) for a particular mRNA.

**Figure 2.**
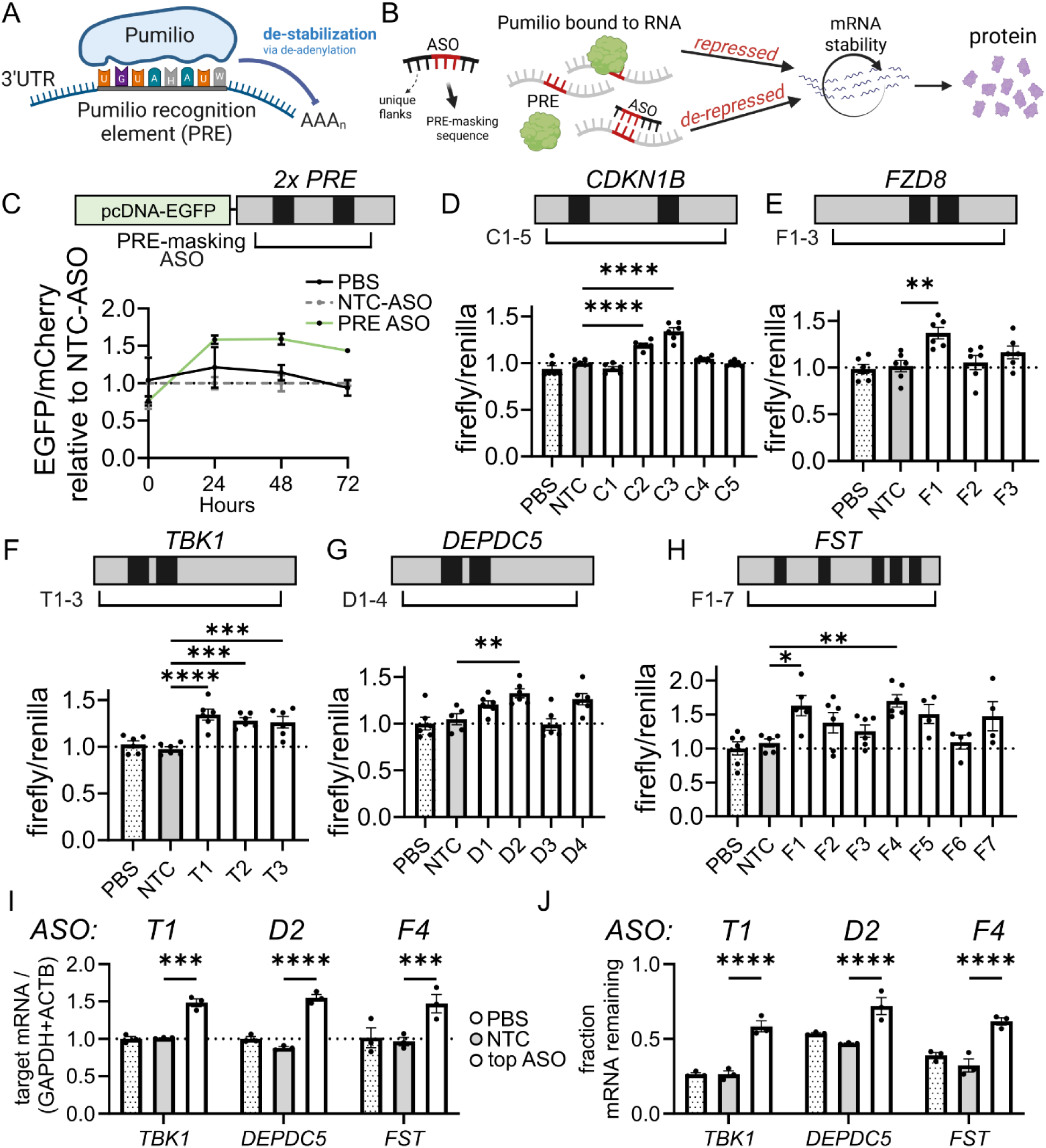
Steric blocking at Pumilio binding sites using antisense oligonucleotides (ASOs) upregulates protein and mRNA. **(A)** Schematic depicting repression of PRE sites by Pumilio proteins. **(B)** Graphic depicting the use of gene-specific antisense oligonucleotides (ASOs) to neutralize PREs for increasing mRNA stability to amplify protein. **(C)** Graphic depicting a fluorescent EGFP reporter flanked by a synthetic 3’UTR that contains two PRE sites, marked by black boxes. Relative fluorescence of EGFP to mCherry control after co-transfection of plasmid reporters with 100nM of specific PRE-masking ASO. Fluorescent readouts were initiated immediately after transfection and continued for 72 hours (n=3 biological replicates per time point; Two-way ANOVA with Dunnett’s post-hoc test, 72hr: p<0.004). (**D, E**) Dual luciferase assays and gene specific ASOs were developed for (D) *CDKN1B* and (E) *FZD8*. The positions of the PRE sites are indicated by black boxes, and gene-specific ASOs by black bars. Dual luciferase plasmids were co-transfected with gene-corresponding ASO, and luminescence was measured after 48 hours (n=5-6 biological replicates; ANOVA with Dunnett’s post-hoc test, *CDKN1B*: p<0.001, *FZD8*: p=0.002). **(F-H)** Dual luciferase assays and gene-specific ASOs were developed for (F) *TBK1*, (G) *DEPDC5*, and (H) *FST*. Plasmids and 100 nM ASOs were delivered to HEK293 cells and luminescence was measured after 48 hours (ANOVA with Dunnett’s post-hoc tests, p<0.05 for each). **(I)** The top ASO candidate (T1, D2, or F4) identified from luciferase assays was delivered to HEK293 cells, and the abundance of the target mRNA was measured by RT-qPCR (n=3 biological replicates; Two-way ANOVA with Dunnett’s post-hoc test, *TBK1*: p<0.0001, *DEPDC5*: p<0.0001, *FST*: p<0.001). **(J)** Cells were transfected with top ASO for 48 hours, then incubated in Actinomycin for 0 hours or 6 hours. The fraction of mRNA from time 0 hours remaining at 6 hours is plotted (n=3 biological replicates per time point; Two-way ANOVA with Dunnett’s post-hoc test, p<0.0001). *p<0.05, **p<0.01, ***p<0.001, ****p<0.0001.

To provide a proof-of-principle for PRE-masking as a strategy for protein upregulation, we utilized fluorescent and luciferase-based reporter systems. We developed a fluorescent EGFP reporter that is flanked by a synthetic 3’UTR that contains two instances of a PRE **(Figure 2C)** and designed reporter-specific ASOs that target individual PRE segments ^27^. Transfecting an empty vector (PBS), non-targeting control (NTC) ASO, or a reporter-specific PRE-targeting ASO revealed that ASOs can increase fluorescent protein production. Protein upregulation by 1.5-fold occurred within 24 hours of ASO treatment and was sustained through 72 hours of incubation (Two-way ANOVA with Dunnett’s post-hoc test, 72hr: p<0.004) **(Figure 2C).**

To demonstrate that this effect is not restricted to *EGFP* mRNA, we implemented dual luciferase reporter systems to extend this approach to 3’UTRs from endogenous genes that contain PREs that are potentially regulated by PUMs. Two well-established targets for PUM proteins are *FZD8* by mRNA destabilization and *CDKN1B* by translation repression. Both *FZD8* and *CDKN1B* have two PREs each (**Figure 2D,E**). We designed ASOs blocking these sites and delivered these ASOs alongside luciferase plasmids encoding the 3’UTR of *FZD8* and *CDKN1B* respectively. Gene-specific ASOs upregulated both 3’UTR reporters by 1.34-fold for CDKN1B-ASO C3 (ANOVA with Dunnett’s post-hoc test, p<0.001) and 1.35-fold for FZD8-ASO F1 (ANOVA with Dunnett’s post-hoc test, p=0.002) (**Figure 2D,E**).

We next focused on three of the disease-associated transcripts repressed by PUM proteins (*TBK1, DEPDC5*, and *FST)* and developed transcript-specific ASOs for each gene. We prepared dual luciferase plasmids for each 3’UTR and delivered each plasmid together with each of its respective ASOs. We found at least one ASO per gene that was able to increase reporter expression by between 1.3- to 1.6-fold (ANOVA with Dunnett’s post-hoc test, p<0.05) (**Figure 2F-H**). Using the top ASO for each gene, we demonstrated that this strategy can also upregulate endogenous gene mRNA for *TBK1* (Two-way ANOVA with Dunnett’s post-hoc test, p<0.0001), *DEPDC5* (p<0.0001), and *FST* (p<0.001) (**Figure 2I**). To confirm a stabilizing mechanism for each ASO, we delivered top ASOs for *TBK1* (T1), *DEPDC5* (D2), or *FST* (F4) to HEK293 cells, then incubated them with transcription inhibitor Actinomycin D for 0 or 6 hours. We calculated RNA stability as the percentage mRNA remaining after the 6 hours incubation compared to the 0 hour baseline. We found that T1, D2, and F4 improved the stability of their respective gene targets (each p<0.0001) (**Figure 2J**). Thus, PRE blocking ASOs can enhance RNA stability in a specific way for multiple genes.

### Pumilio proteins repress TBK1 levels through mRNA de-stabilization

To further demonstrate the mechanism of action, specificity, and applicability of PRE-masking ASOs, we focused on the regulation of TANK-binding kinase 1 (*TBK1*). Haploinsufficiency of *TBK1* is associated with the neurodegenerative disorder amyotrophic lateral sclerosis ^2^. The 3’UTR of *TBK1* contains 2 PREs (**Figure 3A**), and PUM proteins strongly bind *TBK1* mRNA (**Figure 1C**). To evaluate how Pumilio proteins repress TBK1 expression, we depleted PUM proteins by siRNA in HEK293 cells. In addition to mRNA upregulation (**Figure 1D**), we found that depleting PUM proteins increases TBK1 protein by 1.9-fold (t-test, p=0.007) (**Figure 3B**).

**Figure 3.**
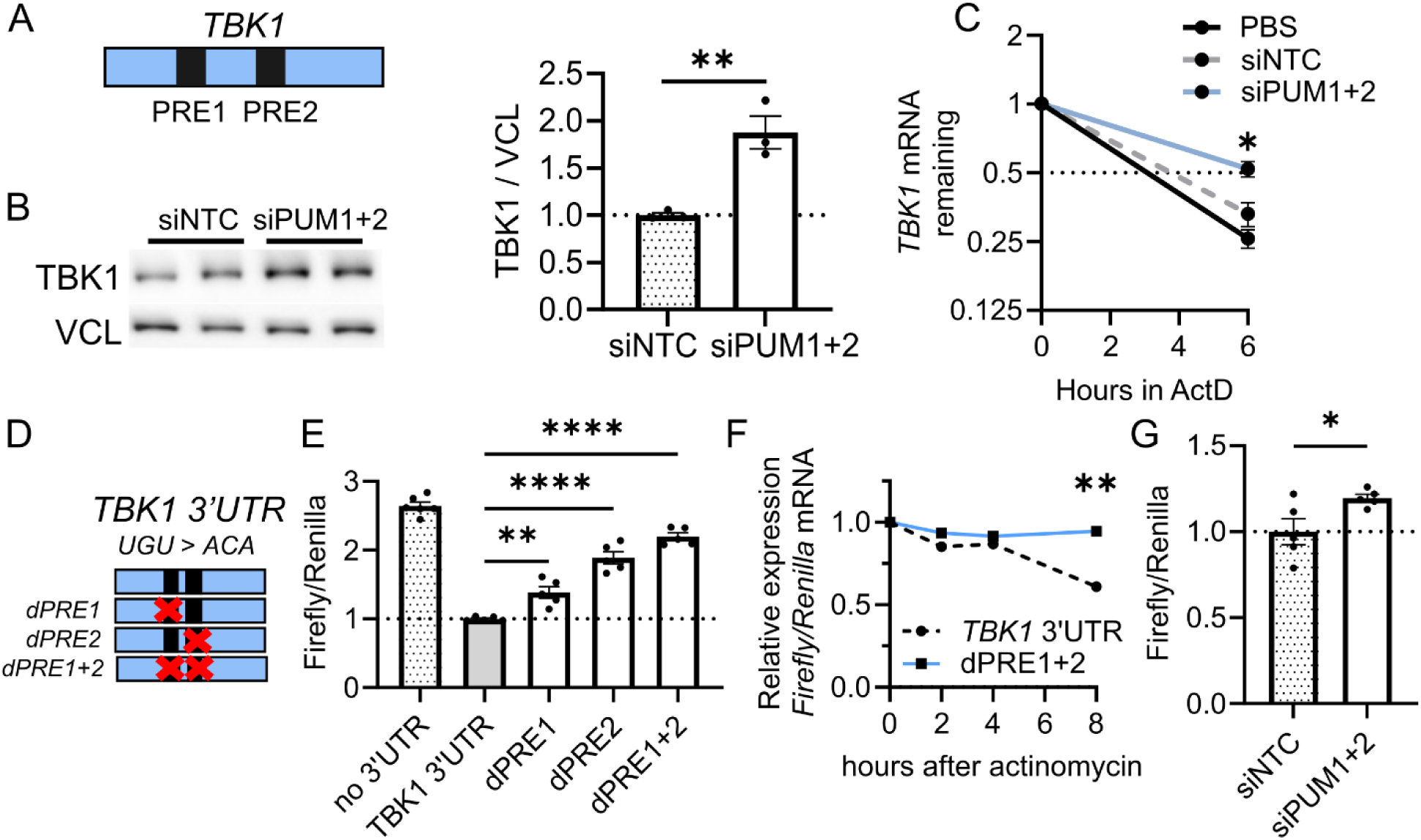
TANK-binding kinase 1 (*TBK1*) is de-stabilized by Pumilio proteins at its recognition elements. **(A)** Depiction of the 3’UTR of *TBK1* with annotated PRE sites by black boxes. **(B)** Western blot and quantification of densitometry after depleting PUM proteins using 100 nM siRNA for 72 hours in HEK293. Vinculin (VCL) was used as a loading control (n=3 biological replicates; t-test, p=0.007). **(C)** After depleting PUM proteins with siRNA, Actinomycin D was delivered for 0 or 6 hours and RNA was harvested. The change in mRNA abundance of *TBK1* from time 0 to time 6 hours is shown (n=3 biological replicates per time point; t-test, p=0.02). **(D)** Strategy to mutagenize PRE sites on *TBK1* 3’UTR dual luciferase assay by converting *UGU* of PRE to *ACA*. **(E)** Relative luminescence from native or mutagenized dual luciferase assays (n=5-6 biological replicates; ANOVA with Dunnett’s post-hoc test, dPRE1: p=0.019; dPRE2: p<0.0001; dPRE1+2: p<0.0001). **(F)** Change in mRNA abundance relative to time 0 after delivering 2xPRE mutagenized plasmid or non-mutagenized control for 48 hours, followed by Actinomycin D for 0, 2, 4, or 8 hours (n=3 biological replicates per time point; Two-way ANOVA, native versus dPRE: p=0.002). **(G)** Relative luminescence after depleting PUM proteins using siRNA for 48 hours (n=6 biological replicates; t-test, p=0.039). *p<0.05, **p<0.01, ****p<0.0001.

Because depleting PUM proteins increased *TBK1* mRNA abundance, we first hypothesized that PUM repression on TBK1 protein expression occurs through mRNA destabilization. To test this possibility, we incubated HEK293 cells with Actinomycin D for 0 or 6 hours after treating with PUM siRNAs. We found that depleting PUMs increased the fraction of *TBK1* mRNA remaining after 6 hours, confirming that PUMs destabilize *TBK1* mRNA (t-test, p=0.02) (**Figure 3C**). To test whether this repression is indeed occurring through PREs on the 3’UTR of *TBK1*, we mutagenized the 3’UTR luciferase reporter at each PRE site and examined protein expression and mRNA stability (**Figure 3D**). We found that adding the full 3’UTR of *TBK1* represses relative luminescence by 2.6-fold. Further, mutagenesis of site 1 increased reporter expression by 1.4-fold, site 2 by 1.9-fold, and loss of both sites elevated reporter levels by 2.2-fold (ANOVA with Dunnett’s post-hoc test, dPRE1: p=0.019; dPRE2: p<0.0001; dPRE1+2: p<0.0001) (**Figure 3E**). In addition, the mutagenized reporter achieved greater mRNA stability after incubating with Actinomycin D for 0-8 hours (Two-way ANOVA, p=0.002) (**Figure 3F**). Finally, depleting PUM proteins increased luciferase expression (t-test, p=0.039) **(Figure 3G)**. These results demonstrate that TBK1 is repressed by PUM proteins and its 3’UTR PREs.

### Optimization of ASO chemistries can increase mRNA upregulation by TBK1-ASOs

ASO chemistries influence ASO function ^28–30^. For example, locked-nucleic acid (LNA) bases or morpholino (MO) backbones offer improved potency and structural rigidity, whereas 2’-methoxyethyl (MOE) bases afford higher safety ^31^. In order to find the most appropriate chemistries for mRNA upregulation, we tested how ASO chemistries affect mRNA upregulation. We optimized ASO chemistries with a fixed sequence from T1 by testing the ratios and positions of LNA versus 2’-MOE bases. We found that ASOs containing two or more LNA bases increased *TBK1* mRNA by >50% consistently (ANOVA with Dunnett’s test, p<0.05) **(Figure S1A)**, which was superior to all MOE sequences at the dose tested. Similarly, configuring the sequence with a fully MO backbone resulted in mRNA increase by 1.8-fold at the highest dose tested (t-test, p<0.01) **(Figure S1B).** Therefore, for further studies, we used a TBK1-ASO containing 18 MOE and 2 LNA bases.

### PRE-masking ASOs increase TBK1 levels through mRNA stabilization

We expected that ASOs targeting the PREs on *TBK1* mRNA operate through eliminating PUM-induced destabilization. To test this possibility, we developed a stable 3’UTR reporter HEK293 cell line that expressed, under doxycycline control, EGFP flanked by the 3’UTR of *TBK1* (**Figure 4A**). We examined whether TBK1-ASO can increase TBK1 and EGFP protein. Indeed, the ASO upregulated each by about 1.7-fold for TBK1 and 2.9-fold for EGFP (ANOVA with Dunnett’s post-hoc test, TBK1: p<0.001, EGFP: p=0.005) (**Figure 4B-D**). Then, we examined whether TBK1-ASOs can increase mRNA abundance of *TBK1* and *EGFP* in a dose-dependent manner. We found a dose-dependent escalation in *TBK1* and *EGFP* mRNA with a >1.8-fold increase in *TBK1* and 1.6-fold increase in *EGFP* at the highest dose tested versus NTC-ASO (t-test, TBK1: p=0.006, EGFP: p<0.05) (**Figure 4E**). To more thoroughly profile mRNA-stabilization by ASOs, we examined *TBK1* and *EGFP* mRNA after ASO and Actinomycin D treatment. ASOs substantially extended mRNA half-life for each gene after treating with Actinomycin D for 0-12 hours (Two-way ANOVA, *TBK1*: p<0.0001, *EGFP*: p<0.0001) (**Figure 4F, G**). Thus, we have now demonstrated using two reporter systems and endogenous TBK1 itself that ASOs operate through targeting the 3’UTR of *TBK1* to increase mRNA and protein expression.

**Figure 4.**
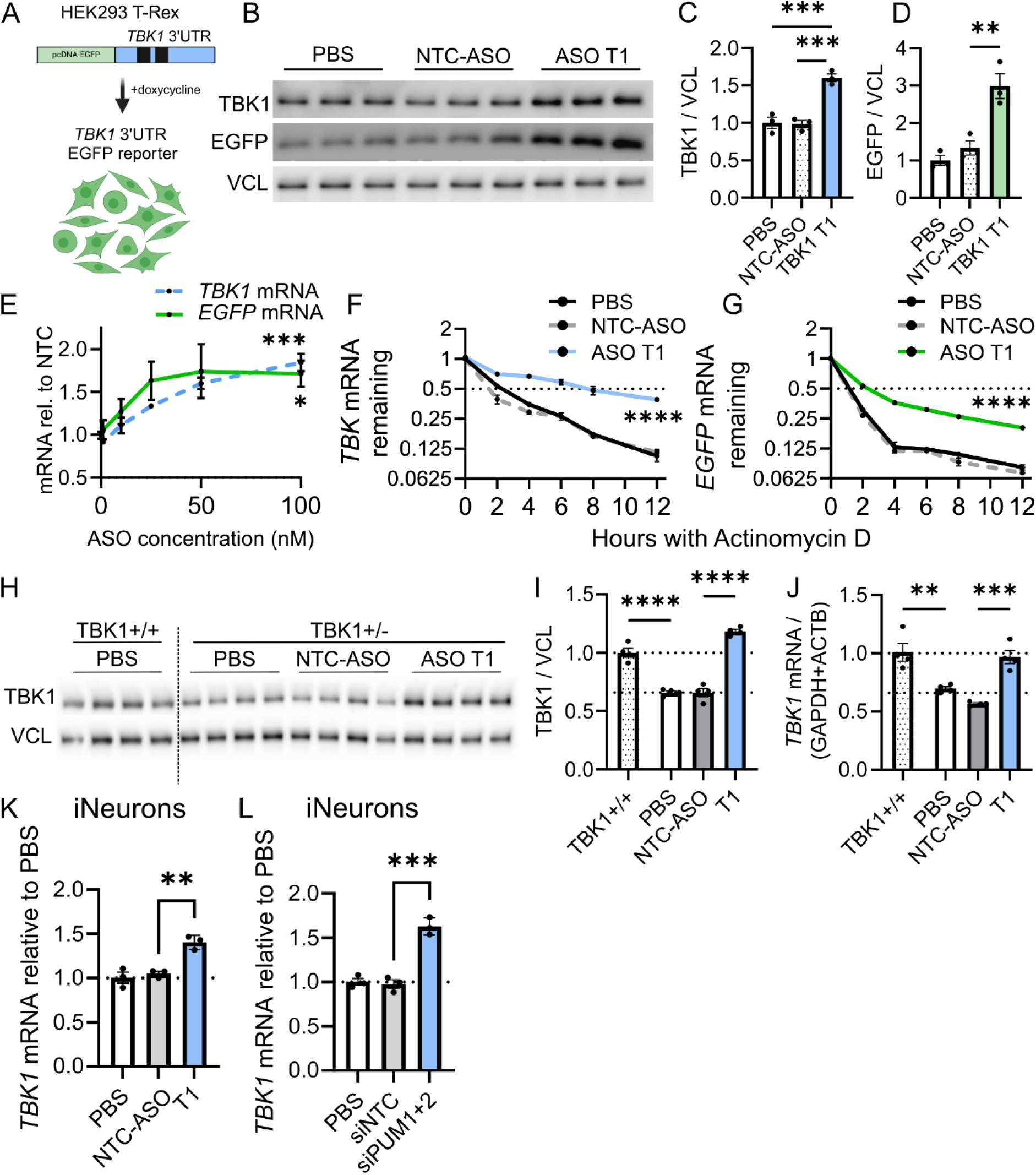
Masking PREs on *TBK1* using ASOs stabilizes *TBK1* mRNA and increases TBK1 protein. **(A)** Graphic depicting reporter HEK293 cell line (T-Rex) that stably expresses EGFP 3’UTR *TBK1* construct under doxycycline control. **(B)** Western blot depicting TBK1 and EGFP protein, alongside vinculin (VCL) loading control, after transfecting HEK293 with 50 nM TBK1-ASO T1 for 72 hours. **(C, D)** Densitometry calculation of TBK1 and EGFP protein relative to VCL loading control. (n=3 biological replicates; t-test, TBK1: p=0.006, EGFP: p<0.01). **(E)** *TBK1* and *EGFP* mRNA abundance with increasing doses of TBK1-ASO T1 after 48 hours (n=3 biological replicates per concentration; t-test, *TBK1*: p=0.006, *EGFP*: p<0.05). **(F-G)** Change in *TBK1* or *EGFP* mRNA after delivering TBK1-ASO T1 for 48 hours to HEK293 cells, then incubated with Actinomycin D for 0, 2, 4, 8, or 12 hours. Half-life was calculated based on change in mRNA abundance over incubation time (n=3 biological replicates per time point; Two-way ANOVA, *TBK1*: p<0.0001, *EGFP*: p<0.0001). **(H)** 100 nM of TBK1-ASO T1 was delivered to TBK1-ALS participant-derived fibroblasts and protein was collected after 72 hours. Western blot depicting TBK1 abundance after treatment with PBS, NTC-ASO, or TBK1-ASO. **(I)** Densitometry of TBK1/VCL after ASO treatment in TBK1+/- fibroblasts (n=4 biological replicates; ANOVA with Dunnett’s post-hoc test, p<0.0001). **(J)** TBK1-ALS participant-derived fibroblasts were treated with TBK1-ASO T1 for 48 hours and mRNA was analyzed by RT-qPCR. Abundance of *TBK1* mRNA was measured (n=4 biological replicates; ANOVA with Dunnett’s post-hoc test, p=0.0004). **(K)** iPSC-derived iNeurons were grown until DIV8, then transfected with 100 nM TBK1-ASO T1 for 72 hours. mRNA was then collected and quantified (n=3 biological replicates; ANOVA with Dunnett’s post-hoc, p<0.01). **(L)** iNeurons were transfected at DIV8 with siRNA against PUM1 and PUM2. *TBK1* mRNA was quantified after 48 hours (n=3 biological replicates; ANOVA with Dunnett’s post-hoc, p<0.001). **p<0.01, ***p<0.001, ****p<0.0001.

### ASOs targeting PREs on *TBK1* can restore protein levels in patient cells and neurons

Using fibroblasts derived from individuals carrying haploinsufficiency mutations in *TBK1*, we examined whether TBK1-ASO can upregulate TBK1 abundance. *TBK1* mutant fibroblasts expressed ∼60% levels of TBK1 mRNA and protein compared to control fibroblast levels, and ASO treatment fully restored TBK1 protein by 1.79-fold (ANOVA with Dunnett’s post-hoc test, p<0.0001) (**Figure 4H-I**) and *TBK1* mRNA by 1.48-fold over NTC-ASO (ANOVA with Dunnett’s post-hoc test, p=0.0004), supporting the translational potential of this strategy (**Figure 4J**). To test in neuronal model systems, we examined whether TBK1-ASO increases *TBK1* mRNA in iPSC-derived iNeurons. ASO treatment increased *TBK1* mRNA by ∼1.4-fold in iNeurons (ANOVA, Dunnett’s test, p<0.01) **(Figure 4K)**. This is consistent with the observation that depleting PUM proteins in iNeurons also upregulates *TBK1* mRNA >1.6-fold (ANOVA, Dunnett’s test, p<0.001) (**Figure 4L**).

### ASOs that mask PREs on *TBK1* are specific

ASOs of 18-24 base pairs that target on or near PRE motifs (8 base pairs) have some potential to bind PREs in other genes. Therefore, we performed two complementary approaches to test the specificity of PRE-masking ASOs for TBK1 (**Figure 5A**).

**Figure 5.**
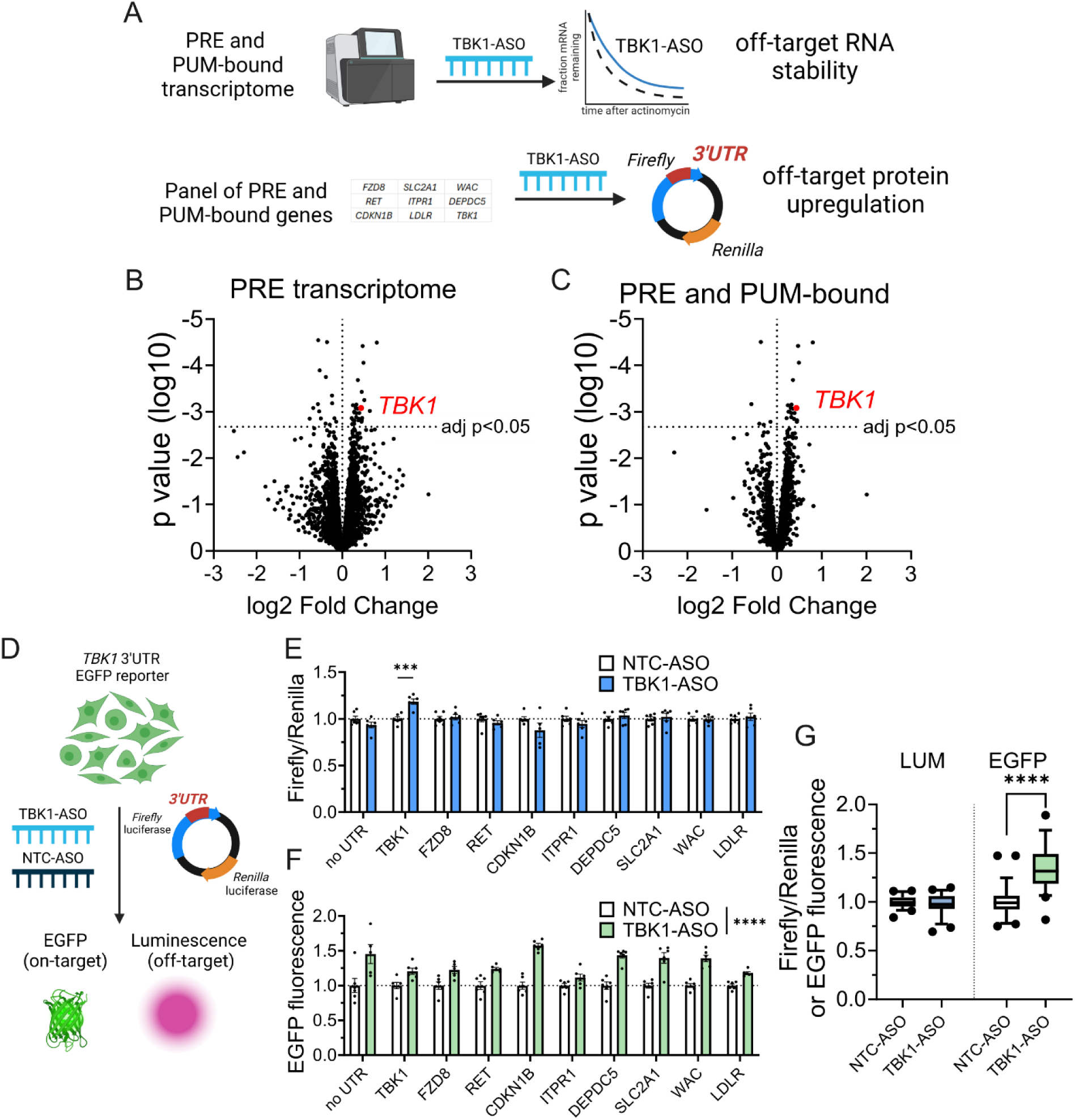
PRE-targeted ASOs disrupt targeted transcripts with high specificity. **(A)** Schematic of two strategies to evaluate specificity by profiling PRE-containing and/or PUM-bound transcriptome or using dual luciferase assays of other PRE containing genes. **(B-D)** TBK1-ASO T1 was delivered to HEK293 cells for 48 hours, then RNA was collected for RNA-sequencing analysis. (B) Volcano plot depicting log2FC of gene targets, after subsetting to genes that contain PRE motifs (5444). (C) Volcano plot depicting log2FC of transcriptome, after subsetting only genes that both contain PRE motifs and are bound (2370). (D) Schematic of testing ASO specificity using 3’UTR fluorescent reporters in HEK293 3’UTR TBK1 reporter cell line. TBK1-ASO T1 was delivered to stable reporter cell line alongside one of ten 3’UTR dual luciferase reporters, including TBK1. For each condition, EGFP and luminescence was recorded after 48-hour incubation. **(E)** Relative luminescence after delivery of non-targeting control ASO or TBK1-ASO T1 on ten 3’UTR reporters (n=5-6 biological replicates Two-way ANOVA with Dunnett’s post-hoc test, TBK1: p<0.001). **(F)** EGFP fluorescence after delivery of non-targeting control ASO or TBK1-ASO T1 on ten 3’UTR reporters (n=5-6 biological replicates Two-way ANOVA, p<0.0001). **(G)** Relative luminescence or EGFP fluorescence is depicted for NTC-ASO or TBK1-ASO after averaging each condition. For this analysis, the TBK1 3’UTR luciferase reporter was excluded from the luminescence measure (t-test, p<0.0001). ***p<0.001, ****p<0.0001.

In the first approach, we examined all genes differentially expressed with TBK1-ASO treatment using an unbiased RNA-sequencing approach. To isolate RNA changes related to ASO versus TBK1 upregulation, we collected RNA in HEK293 after a 48-hour ASO treatment. Further, we sought to increase the power to detect off-target effects by examining only transcripts that (1) contain PREs or (2) both contain PREs and are bound by PUMs ^10,26^. We identified 36 genes, including *TBK1* that were differentially expressed in either group (19 genes that were significant when considering only PREs, 24 for bound by PUMs, and 23 that were significant when considering both criteria) (**Figure 5B-C** and **Table S4-5**) – either because they are also bound and stabilized by our ASO (‘direct’ off-target effects) or because they happen to be upregulated downstream to TBK1 elevation (indirect effects). To distinguish these possibilities, we first evaluated the alignment of our ASO to the UTRs of these 30 genes via a low-stringency BLAST. The top three genes *SLC5A3*, *HCG11*, and *EXO1* had 3, 9, and 8 mismatches, respectively. The presence of two or more mismatches in a sequence is sufficient to eliminate ASO activity at that site ^32,33^. Thus, it appears these likely do not represent any worrisome off-target effects, but rather normal downstream and TBK1-driven upregulations. To further confirm this, we repeated the ASO exposure, but with Actinomycin D, followed by RT-qPCR to assess stability, as direct off-target effects would influence stability, while indirect (likely transcriptional effects) would not. Validating the RNA-seq results, we found that TBK1-ASO alters gene expression of these targets (ANOVA with Dunnett’s post-hoc test, *TBK1*: p=0.002, *SLC5A3*, p=0.03, *HCG11*, p=0.03, *EXO1*, p>0.05) (**Figure S2A**), yet only *TBK1* had altered mRNA stability after Actinomycin D incubation (t-test, p>0.05 for *SLC5A3, HCG11,* and *EXO1*) (**Figure S2B**). Overall, ASOs targeting the PRE on *TBK1* appear to have limited off-target effects due to shared PRE motifs.

As a complementary approach, we examined whether ASOs influence protein abundance of potential off-target PRE-containing genes. To test this, we developed and implemented luciferase assays for 10 genes. We included three established Pumilio targets *FZD8*, *CDKN1B*, and *RET*, as well as previously used luciferase reporters for above PUM-regulated genes. To validate delivery and target engagement of our ASOs, we transiently transfected each luciferase reporter to the EGFP 3’UTR reporter cells so we could simultaneously confirm on-target effects of ASO on the *TBK1* 3’UTR (**Figure 5E**) by EGFP, and assess any off-target effects by luciferase, owing to its higher sensitivity. We confirmed EGFP fluorescence was upregulated by TBK1-ASO (Two-way ANOVA, p<0.0001) in all conditions **(Figure 5F)**, while luminescence of potential off-target 3’UTR reporters demonstrated that only TBK1 luciferase reporter levels were elevated with ASO (Two-way ANOVA with Dunnett’s post-hoc test, TBK1: p<0.001) **(Figure 5G)**. On average, TBK1-ASO did not change off-target luminescence (t-test, p>0.1) but increased average fluorescence by 1.36-fold relative to NTC-ASO (t-test, p<0.0001) **(Figure 5H).** These data are consistent with simultaneous on-target engagement at TBK1 without off-target effects, at the protein level.

Collectively, these complementary approaches illustrate that effects of PRE-targeted ASO for TBK1 are highly specific to TBK1 at both the RNA and protein levels.

### ASOs targeting PREs in *DEPDC5* and *CDKN1B* influence cellular state

We anticipated that upregulating target genes with this strategy would yield functional outcomes. To test this, we focused on ASOs targeting the two haploinsufficiency-associated genes *DEPDC5* (epilepsy) and *CDKN1B/p27Kip* (leukemia) that have well-characterized cellular readouts. DEPDC5, part of the GATOR1 complex, inhibits mTORC1, a key autophagy regulator ^22,34^. CDKN1B is a cell cycle inhibitor (p27Kip) and haploinsufficient tumor suppressor, blocking cell cycle progression from G1 to S phase ^35,36^. Augmenting its expression should delay S phase entry.

To demonstrate DEPDC5 upregulation with ASOs impacting mTOR signaling, we tested the potential of DEPDC5-ASO D2 to activate autophagy by examining autophagy substrate accumulation after employing Bafilomycin A (BafA), an autophagolysosomal fusion inhibitor.

After treating HEK293 cells for 48 hours with ASO D2, we incubated cells with Bafilomycin A for 6 hours and examined autophagy regulators. We observed that D2 trends towards suppressing p62 (ANOVA with Dunnett’s post-hoc test, p=0.06) (**Figure S3A,B**) and increasing LC3-II abundance (**Figure S3A,C**). In Bafilomycin-treated cells, D2 decreased p62 (ANOVA with Dunnett’s post-hoc test, p=0.01) (**Figure 6A,B**) and increased LC3-II (ANOVA with Dunnett’s post-hoc test, p=0.02) (**Figure 6A,C**). These findings indicate more active degradation and strong autophagy flux, consistent with DEPDC5 influencing autophagy.

**Figure 6.**
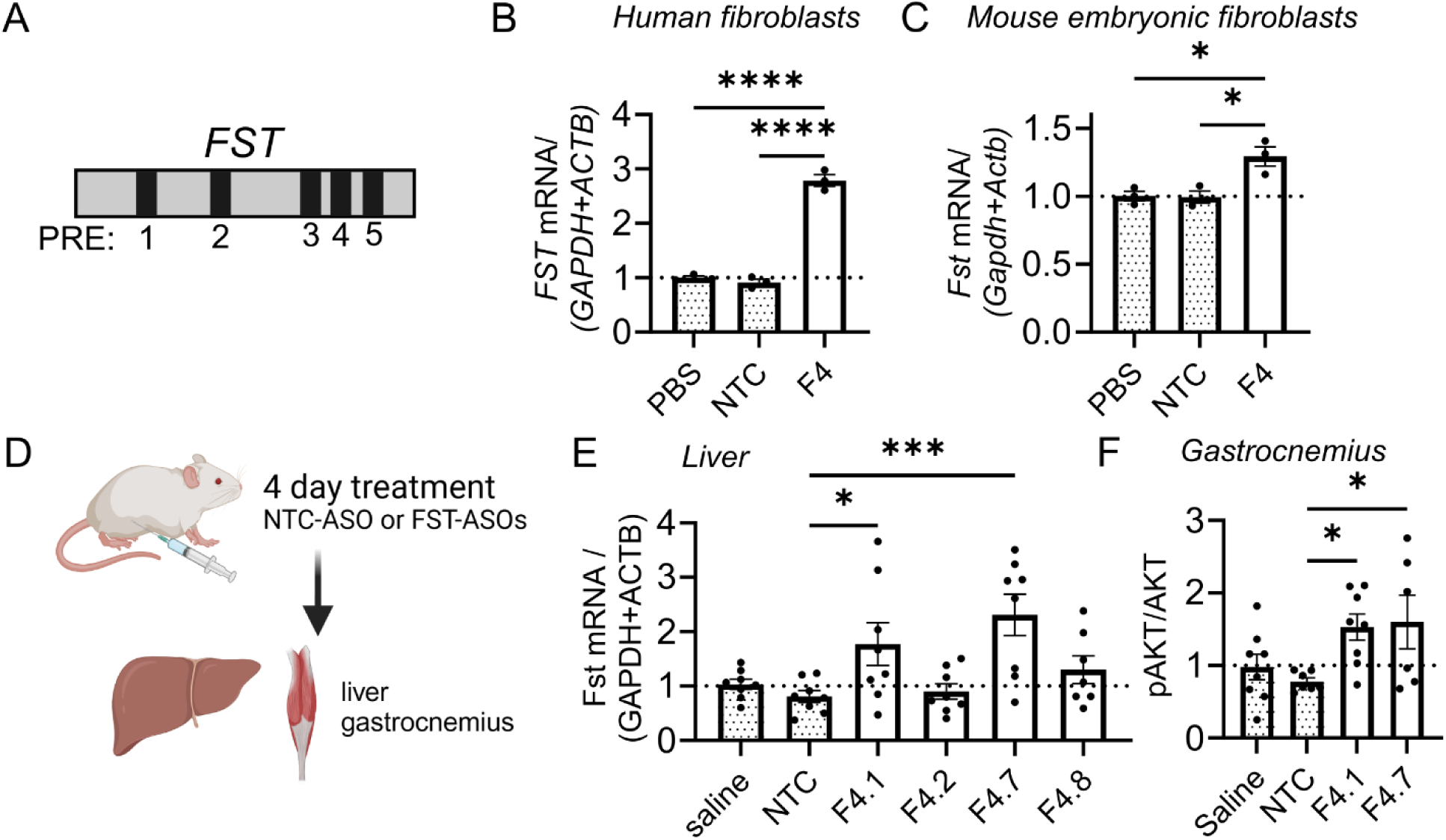
Conserved PRE-masking ASOs increase gene expression *in vivo*. **(A)** Depiction of the 3’UTR of *FST* with annotated PRE sites by black bars. **(B,C)** Quantification of relative *FST* mRNA after delivery of 50nM FST-ASOs to (B) human fibroblasts ((n=3 biological replicates; ANOVA, Dunnett’s test, p<0.0001) or (C) mouse embryonic fibroblasts for 48 hours (n=3 biological replicates; ANOVA, Dunnett’s test, p=0.01). **(D)** Schematic for delivery of FST-ASO to the mouse by intraperitoneal injection. After a 4-day treatment with ASO, liver tissue and gastrocnemius muscle were collected. **(E)** Quantification of relative *Fst* mRNA in the right lobe of the liver after intraperitoneal delivery of 25mg/kg FST-ASOs, saline, or NTC-ASO for 4 days (n=6-8 biological replicates; ANOVA with Dunnett’s test, F4.1: p<0.05, F4.7: p<0.001). **(F)** The proportion of pAKT/AKT was measured in the gastrocnemius muscle after treatment with FST-ASO 4.1 or 4.7 by immunoblotting. Densitometry analysis is depicted (n=6-8 biological replicates; ANOVA, Dunnett’s test: FST 4.1 p<0.05, FST 4.7 p<0.05). *p<0.05, ***p<0.001, ****p<0.0001.

We also tested if CDKN1B ASO C3, which most strongly upregulated the *CDKN1B* 3’UTR reporter, could increase cell cycle arrest. Using a cell cycle reporter (Fucci(SA)5) (**Figure S3D**) ^37^ stably integrated in HeLa cells, we delivered 25nM ASO C3 for 48 hours, then performed flow cytometry to quantify cell cycle state. C3 increased the proportion of cells at the G1/S checkpoint compared to a NTC-ASO control (t-test, p=0.004) (**Figure S3F**). This occurred alongside a decrease in cells in G2/S/M phases (t-test, p=0.002) and trended increase in cells in G1 (t-test, p=0.07) (**Figure S3G,H**). These findings indicate the ability for CDKN1B ASOs to influence cell cycle progression.

### mRNA-stabilizing ASOs targeting PREs are active *in vivo*

Finally, we sought to evaluate whether PRE-targeting ASOs can increase protein abundance in a whole organism using a mouse model. While the PREs in *TBK1* are conserved across species, the surrounding gene sequences are sufficiently different to make the human-targeted ASOs inactive against the mouse *TBK1* sequence. Of the other ASOs we had characterized, *FST* mRNA is strongly repressed by PUM proteins (**Figure 1D**), and the sequences targeted by FST-ASOs are fully conserved across species; therefore, we selected ASOs targeting *FST* for *in vivo* testing (**Figure 6A**). *FST* encodes follistatin, which is a secretory protein that is expressed ubiquitously and inhibits myostatin signaling in muscle to promote muscle growth.

*FST* is abundantly expressed in human fibroblasts, so we tested if *FST*-targeted ASOs could increase *FST* mRNA *in vitro*. ASOs strongly upregulated *FST* by >2.5-fold in fibroblasts (ANOVA, Dunnett’s test, p<0.0001) (**Figure 6B**). To examine whether these ASOs are also active against mouse *Fst*, we delivered FST-ASOs to mouse embryonic fibroblasts and again achieved upregulation of *Fst* mRNA (ANOVA, Dunnett’s test, p=0.01) (**Figure 6C**). We optimized candidate ASOs, based on ASO F4, to advance to *in vivo* study (**Figure S4**). Since most *Fst* is produced in the liver, we delivered *FST*-targeted ASOs by intraperitoneal injection at 25mg/kg to C57BL/6 naïve mice and harvested liver and muscle after four days (**Figure 6D**). *FST-*targeted ASOs increased *Fst* mRNA by 2 to 2.9-fold in the liver (ANOVA with Dunnett’s test, F4.1: p<0.05, F4.7: p<0.001) (**Figure 6E**).

Follistatin achieves its myoprotective activity by inhibiting myostatin signaling in the muscle, so we examined whether FST-ASOs impact myostatin signaling. Indeed, ASOs 4.1 and 4.7 both augmented pAKT/AKT levels by >1.5-fold (ANOVA, Dunnett’s test: FST 4.1 p<0.05, FST 4.7 p<0.05), which have previously demonstrated upregulation with FST gene therapies (**Figure 6F**) ^38,39^. These results demonstrate that PRE-targeting ASOs can operate *in vivo* to upregulate gene expression and produce functional effects.

## DISCUSSION

In this work, we introduced a novel strategy using 3’UTR-targeted ASOs to amplify protein expression. We focused first on Pumilio recognition elements (PREs), likely applicable to 30-40% of the transcriptome, demonstrating that ASOs can upregulate PRE-containing 3’UTR reporters for several disease-associated genes. We established the ALS-associated gene *TBK1* as a novel PUM target that can be de-repressed using ASOs targeting its PREs. Using TBK1-ASOs as a case example, we demonstrated the specificity of PRE-targeting ASOs and their ability to rescue gene expression in neurons and participant-derived fibroblasts. Using *FST* PRE-targeting ASOs, we further showed the ability of PRE-targeting ASOs to upregulate gene expression *in vivo*.

There are over 300-1000 haploinsufficient genes, and the vast majority lack gene-targeted therapeutics ^1^. Here, we identify ASOs that upregulate deficient gene expression or activity for forms of ALS/FTD (*TBK1*), familial epilepsy (*DEPDC5*), and leukemia (*CDKN1B*) ^2,21,22,25^. Given that haploinsufficiency often only entails ∼50% loss of protein, we consider the degree of upregulation by ∼1.7x achieved by TBK1 PRE-targeting ASOs as promising and predict that this degree of increase will provide benefit in TBK1-related ALS/FTD. Just as with TBK1-ASOs, further development of ASOs for other haploinsufficiency disorders could be transformative for these orphan diseases. ASOs that readily upregulate gene expression could circumvent many challenges of other gene upregulation approaches (such as viral gene transfer or direct protein delivery), which can have challenges with safety and dosing. Here, we establish a proof-of-principle, using *TBK1*-related ALS, for an ASO approach that has potential to generalize for many genes of interest.

Gene regulation at the 3’UTR occurs through many unique inhibitory and activating forces, with microRNA binding sites, AU-rich elements (AREs), and PREs representing three canonical post-transcriptional repressors. Similar to our work with PUM here, previous groups have demonstrated that ASOs blocking microRNA sites on selective 3’UTRs can effectively upregulate gene expression and reverse disease phenotypes ^40,41^. However, generalizing a microRNA-based approach is challenged by cell-restricted expression of some microRNAs and the limited control that each microRNA has on any single transcript. Similar challenges are observed with AREs, which can have variable activity at each transcript and cell-specific effects ^42–45^. Beyond these repressive sequences, structural features and mRNA modifications can influence mRNA stability and translation. The rationale-design, regulatory-element informed strategy implemented here may fail to identify these UTR features. Therefore, for a single gene of interest, emerging discovery-based scanning strategies may be able to discover UTR-specific repressive elements in a high-throughput manner.

Based on our luciferase studies of UTR mutations, a theoretical 2.2-fold upregulation is possible through PRE de-repression, which is not yet achievable by our existing ASO candidates. Thus, our current ASOs might be augmented through improved ASO sequences or chemistries or combinatorial treatment with microRNA-blocking ASOs. We predict that targeting regulatory regions on 3’UTRs controlled by multiple repressors, such as adjacent binding of Pumilio proteins and microRNAs, are highly attractive targets for this strategy. Indeed, Pumilio has an established role in coordinating microRNA activity at 3’UTRs through mRNA structural remodeling, such as in controlling CDKN1B expression ^46,47^. High AU content, such as in AREs, around Pumilio and microRNA recognition sites generally supports their repressive activity ^10^. Additionally, we expect combinatorial ASO treatment for multiple elements in the same UTR could potentially provide even greater activation in amenable cases.

While dozens to hundreds of disorders are related to haploinsufficiency, protective signaling pathways are under investigation across human conditions. Using follistatin-upregulating ASOs as a proof-of-principle, we introduced the potential for applying this strategy beyond haploinsufficiency syndromes. These findings indicate that blocking the tonic repression by PREs on a transcript can be used to achieve supra-normal gene upregulation. Follistatin is a secreted glycoprotein that can promote muscle growth through antagonism of myostatin signaling ^24,48^. Myostatin antagonism, such as with FST gene therapy, has faced challenges in clinical trials for muscular disorders thus far but is an approach that remains actively investigated ^49^. However, FST-ASOs provide the possibility of translating this approach to other protective proteins and signaling pathways. For example, we expect this approach to be applicable to upregulate growth factors such as vascular endothelial growth factor (VEGFA) or glial-derived neurotrophic factor (GDNF) that might provide neuroprotection against aging or neurodegenerative disorders ^50,51^.

There are some limitations to our study. For example, we focused on introducing ASO suppression of PREs as a concept, but we acknowledge that we did not advance TBK1-ASOs to advanced pre-clinical or humanized mouse models, which would further establish strong therapeutic relevance. Nevertheless, we consider the ASOs we show here as promising, strong therapeutic candidates. We have established here that PUM sites are regulatory regions that can be manipulated and have selected and tested these empirically. However, to fully take advantage of this approach, we need to develop a high throughput system to examine PREs and other potential regulatory regions in the 3 ‘UTR that could be applied to any gene, an approach that would be particularly helpful for genes with long 3’UTRs. We expect future studies to define this new class of gene-augmenting drugs and how they can be leveraged in pre-clinical conditions, such as related to TBK1 haploinsufficiency.

Overall, we establish a generalizable approach for gene upregulation that can be applied to restore dozens of haploinsufficiency genes or augment protective signaling pathways. Our initial focus on TBK1-upregulating ASOs has already introduced promising leads on a potential therapy for TBK1 haploinsufficient ALS/FTD. This strategy opens new avenues for treating haploinsufficiency disorders and enhancing protective protein signaling across various diseases.

## METHODS

### Study design

This study was designed to establish a novel ASO-based approach to enhance protein expression by stabilizing mRNA transcripts through targeted masking of repressive elements within the 3′UTR. Outcomes were measured by assessing mRNA stability, protein expression levels, and functional restoration in disease-relevant cellular and animal models. We focused on PRE-masking ASOs for *TBK1* that restored TBK1 protein expression through mRNA stabilization in fibroblasts derived from individuals with TBK1-ALS/FTD. We further evaluated efficacy of PRE-masking ASOs *in vivo* for a myoprotective gene *FST* and observed effective upregulation in target tissues.

For animal studies, mice were randomly assigned to treatment groups and received ASOs via intraperitoneal (IP) injection. No animals were excluded from analysis. Age-matched littermates were used as controls across all *in vivo* experiments. All animal procedures were carried out in compliance with national and institutional ethical guidelines, with prior approval from the Institutional Animal Care and Use Committee at Washington University in St. Louis.

All cellular experiments were conducted with at least 3 biological replicates. Animal studies were performed with at least 5 biological replicates. Quantification of mRNA by RT-qPCR was performed with 2 technical replicates.

### Cell culture

Human embryonic kidney 293 (HEK293) cells were obtained from the American Type Culture Collection (CRL-1573; ATCC, Manassas, VA, USA). HeLa cells were obtained from the American Type Culture Collection (CCL2; ATCC, Manassas, VA, USA). HEK293 and HeLa cells were maintained in DMEM and 10% FBS, containing antibiotics, unless described otherwise.

Stable Flp-In T-REx-HEK293 cells expressing an inducible EGFP *TBK1* 3’UTR reporter were generated using the Flp-In T-REx system (Invitrogen). The EGFP coding sequence followed by the *TBK1* 3’UTR was cloned into the pcDNA5/FRT/TO vector. Flp-In T-REx-HEK293 host cells were co-transfected with the pcDNA5/FRT/TO-EGFP-3’UTR construct and pOG44 Flp recombinase expression plasmid at a 1:9 ratio using Lipofectamine 3000 (Invitrogen) according to the manufacturer’s protocol. Forty-eight hours post-transfection, cells were selected with 100 μg/mL hygromycin B for two weeks. Individual colonies were isolated, expanded, and screened for tetracycline-inducible EGFP expression. EGFP expression was induced with 1 μg/mL tetracycline for 24 hours and verified by fluorescence microscopy and flow cytometry. Positive clones were maintained in DMEM supplemented with 10% tetracycline-free FBS, 100 μg/mL hygromycin B, and 15 μg/mL blasticidin. Induction of EGFP in cell culture was performed by adding doxycycline to final concentration of 1ug/mL in the media without replacement.

iPSC-derived iNeurons with a doxycycline-inducible NGN2 promoter were cultured and differentiated using a modified protocol. On day 0, cells were seeded onto a Matrigel-coated (Corning) plate into Stemflex media (Thermo Fisher Scientific) without antibiotics, supplemented with 10 μM ROCK inhibitor Y-27632 (Tocris). Media was changed daily after the first day. On day 1, the culture was switched to N2B27 media (1:1 mixture of DMEM/F12 and Neurobasal media supplemented with 1% N2, 2% B27, 1% GlutaMAX, and 0.1% β-mercaptoethanol, all from Thermo Fisher Scientific). Cells were maintained in N2B27 media with daily changes until day 5. On day 5, cells were dissociated using Accutase (Stemcell Technologies) and re-plated onto 12-well plates coated with poly-d-lysine (Sigma) in N2B27 media containing 10 μM Y-27632. Neuronal differentiation was induced by adding 2 μg/mL doxycycline to the culture media. On day 8 (DIV8), cells were transfected using Lipofectamine RNAiMAX (Invitrogen) according to the manufacturer’s protocol. Media was changed 24 hours post-transfection. Cells were maintained until DIV10, at which point they were collected for downstream analyses. Throughout the differentiation process, cells were kept at 37°C in a humidified incubator with 5% CO2.

### Plasmid constructs and mutagenesis

The 3’UTRs of interest (*TBK1, DEPDC5, ITPR1, LDLR, FST, SLC2A1, WAC)* were amplified from human genomic DNA. The PCR reactions were performed using Q5 High-Fidelity DNA Polymerase (New England Biolabs) according to the manufacturer’s instructions. The pMirGlo vector (Promega) were digested with XhoI and NotI restriction enzymes (New England Biolabs) for 1 hour at 37°C. The digested vector was dephosphorylated using Antarctic Phosphatase (New England Biolabs) to prevent self-ligation. The PCR products were purified using the Monarch PCR cleanup kit (NEB). The In-Fusion cloning reaction was set up using the In-Fusion Snap Assembly Master Mix (Takara Bio) with a 3:1 molar ratio of insert to vector. The reaction was incubated at 50°C for 15 minutes, then placed on ice. The ligation products were transformed into competent E. coli DH5α cells (Invitrogen) using heat shock at 42°C for 30 seconds and according to manufacturer instructions. Transformed bacteria were plated on LB agar containing 100 μg/mL ampicillin and incubated overnight at 37°C. Individual colonies were screened by colony PCR, and positive clones were confirmed by Sanger sequencing (Genewiz).

Mutations in the PRE sites of TBK1 were introduced using site-directed mutagenesis. Mutagenic primers were designed with Quickchange software. PCR amplification was performed using Phusion High-Fidelity DNA Polymerase (Thermo Fisher Scientific) with 5 ng of template DNA, 0.5 μM of each primer, 200 μM dNTPs, and 1X Phusion HF Buffer in a 50 μL reaction volume. Thermal cycling conditions were: initial denaturation at 98°C for 30 seconds; 19 cycles of 98°C for 10 seconds, 65°C for 30 seconds, and 72°C for 8 minutes; followed by a hold at 4°C. The PCR product was treated with 1 μL DpnI (New England Biolabs) at 37°C for 1 hour to digest the parental DNA template. NEB 5-alpha Competent E. coli cells (New England Biolabs) were transformed with 5 μL of the DpnI-treated PCR product using heat-shock. Transformants were selected on LB agar plates containing appropriate antibiotics. Positive clones were identified by colony PCR and confirmed by Sanger sequencing (Genewiz).

### Cell cycle analysis

A monoclonal HeLa cell line stably integrating doxycycline inducible tFucci(SA)5 ^52^ and rtTA3G was established using the Sleeping Beauty transposon system. Cells were cultured in medium containing 2.5 μg/mL doxycycline. Cells were transfected with ASOs or a control RNA oligonucleotide corresponding to pre-cel-miR-67 at a final concentration of 25nM using Lipofectamine RNAiMAX. 48hrs post transfection cells were harvested, washed twice and resuspended in cell staining buffer (Biolegend) and analyzed using a MACSQuant VYB flow cytometer (Miltenyi). Downstream analysis was performed using FlowJo software. Cells in the G1 phase displayed red fluorescence (mKO2-Cdt1), while cells in S/G2/M phases exhibited green fluorescence (mAG-Geminin). The proportion of cells in each cell cycle phase was quantified based on population gating.

### Dual luciferase assays

Firefly and Renilla luciferase activities were quantified using the Dual-Glo® Luciferase Assay System (Promega) following the manufacturer’s protocol. Cells were seeded in 96-well plates and transfected after 24 hours with the experimental Firefly/Renilla dual luciferase reporter construct using Lipofectamine 3000 (Thermo Fisher Scientific). After 48 hours, cells were lysed in 50 μl Dual-Glo® reagent, incubated for 10 minutes at room temperature, and firefly luciferase activity was measured using a luminometer (Agilent, BioTek Synergy H1). Subsequently, 50 μl of Dual-Glo® Stop & Glo reagent was added to quench firefly luciferase activity and activate Renilla luciferase, followed by a 30-minute incubation and measurement. Results were normalized to Renilla luciferase activity to control for transfection efficiency and cell viability. All experiments were repeated with at least three independent biological replicates. Empty vector and mock-transfected cells served as controls.

### Drug treatments

For transcriptional inhibition, cells were treated with Actinomycin D (ActD, Sigma-Aldrich A9415) at a final concentration of 5 μg/mL for 0-12 hours. At each time point, cells were harvested for RNA extraction and subsequent analysis. The 0-hour time point served as the baseline for normalization. For autophagy inhibition, cells were treated with Bafilomycin A1 (BafA, Sigma-Aldrich B1793) at a final concentration of 100 nM for 4 hours. Control cells were treated with an equivalent volume of DMSO (vehicle).

### ASO designs

All ASOs were designed as indicated and ordered from Integrated DNA Technologies or GeneTools (only morpholinos). All were fully modified 15-25 base pair sequences consisting of 2’-methoxyethyl or locked nucleic acid bases. Backbones used were phosphorothioate or morpholino. ASO sequences and chemistries may be provided upon request and will be disclosed upon publication.

### ASO and siRNA treatments in cells

Treatments using ASOs or siRNA were performed using reverse transfection unless otherwise indicated. HEK293 cells were plated in DMEM at 75% confluency at the time of transfection. ASO or siRNA was complexed with RNAiMAX transfection reagent (Invitrogen) in Opti-MEM Reduced Serum Medium according to manufacturer recommendations, with final ASO or siRNA concentrations ranging from 10-250 nM in culture medium. The RNAiMAX:ASO complexes were incubated at room temperature for 20 minutes before being added dropwise to cells. Transfected cultures were maintained for 48-72 hours at 37°C with 5% CO2, without media replacement. Non-targeting ASO controls and lipofectamine only (PBS) cells serving as negative controls. Cells were harvested for downstream analysis (qPCR, western blot, or functional assays) at experimental endpoints.

### Immunoblotting

Cells were washed once with ice-cold PBS and lysed in RIPA buffer (50 mM Tris-HCl pH 7.4, 150 mM NaCl, 1% NP-40, 0.5% sodium deoxycholate, 0.1% SDS) supplemented with fresh EDTA-free protease inhibitor cocktail (Roche Diagnostics). Lysates were sonicated for 5 minutes at 4°C, followed by centrifugation at 21,000 × g for 15–20 minutes at 4°C to pellet insoluble debris. Supernatants were quantified using the Pierce BCA Protein Assay Kit according to the manufacturer’s protocol, with absorbance measured at 562 nm after a 30-minute incubation at 37°C. Protein concentrations were normalized against a bovine serum albumin (BSA) standard curve. For electrophoresis, 20 µg of protein per sample was mixed with Laemmli buffer, denatured at 95°C for 10 minutes, and resolved on a 4%–20% gradient polyacrylamide gel (Bio-Rad) using Tris-glycine-SDS running buffer. Proteins were transferred to PVDF membranes using a transfer system at constant voltage (400 mA, 1 hour) for downstream immunoblotting.

Membranes were incubated in primary antibodies overnight, then 1 hour incubation in secondary antibody. Membranes were visualized using Clarity ECL reagent (Biorad, 1705061). Primary antibodies used for blotting: VCL at 1:4000 (Sigma, V9131), TBK1 at 1:1000 (Abcam, ab40676), pAKT at 1:1000 (Cell Signaling Technologies, 9271S), AKT at 1:1000 (Cell Signaling Technologies, 9272), EGFP at 1:1000 (Invitrogen MA1-952), p62 at 1:3000 (Abcam, ab56416), and LC3b at 1:1000 (Cell Signaling Technologies, 83506). Secondary antibodies used: anti-Rabbit IgG HRP (Sigma, GENA934) and anti-Mouse IgG HRP (Sigma, GENA931).

### mRNA quantification

RNA was isolated from cell, liver, or muscle tissue using the RNeasy Mini Kit (Qiagen) following the manufacturer’s instructions. Cells or tissue were lysed with 300 μL RLT buffer (Qiagen) in the plate or in Eppendorf tubes. If using cells, the plate was shaken for 10 minutes at room temperature. The sample was then mixed with 1.5 volumes of 100% ethanol and transferred to an RNeasy column for RNA purification with DNase treatment according to the kit protocol. cDNA synthesis was carried out using the High-Capacity cDNA Reverse Transcription Kit (Invitrogen). Quantitative PCR was performed on the QuantStudio 12K Flex Real-Time PCR System using Power SYBR Green Power PCR Master Mix (Thermo Fisher Scientific). Gene expression was quantified using the ΔΔCt method, using *GAPDH* and *ACTB* (human) or *Gapdh* and *Actb* (mouse) as reference genes.

RNA-seq analysis was performed on samples sequenced using an Illumina NovaSeq X Plus platform. Raw data processing, including basecalling and demultiplexing, was conducted using Illumina’s DRAGEN and BCLconvert (v4.2.4) software. Reads were aligned to the Ensembl release 101 primary assembly using STAR (v2.7.9a1), with gene counts determined by Subread:featureCount (v2.0.32) and isoform quantification by Salmon (v1.5.2). Sequencing quality was assessed using RSeQC (v4.0). Data normalization and differential expression analysis were performed using EdgeR and Limma R/Bioconductor packages, with TMM normalization and voomWithQualityWeights transformation. Only genes containing Pumilio recognition elements or that are bound by Pumilio were compared. Benjamini-Hochberg FDR-adjusted p-values ≤ 0.05 based on this subsetted genome were considered significantly differentially expressed.

### Animals

C57BL/6J mice (The Jackson Laboratory, stock no. 000664) were bred and housed at Washington University in St. Louis. Mice were provided with unlimited food and water and kept on a standard 12-hour light/dark cycle. Animal use was conducted in accordance with the National Institutes of Health (NIH) guidelines for animal research, under protocols approved by the Institutional Animal Care and Use Committee of Washington University in St. Louis.

### ASO intraperitoneal injection

All experiments were performed using male C57BL/6J mice. Mice received a single intraperitoneal (IP) injection of saline, non-targeting control (NTC) ASO, or an *FST*-targeting ASO at a dose of 25 mg/kg prepared in sterile saline. Four days after IP injection, mice were perfused with cold PBS and euthanized. The liver and the right and left gastrocnemius muscles were rapidly dissected. Tissues were immediately flash-frozen in liquid nitrogen and subsequently stored at -80°C until further processing and analysis.

### Statistical analysis

For all experiments, the primary comparison was between NTC-ASO control versus any targeting ASOs that were tested. Comparisons between two groups were conducted using student’s t-test. Multiple comparisons were conducted using ANOVA with Dunnett’s post-hoc test, such as in ASO screening experiments. Measurements of mRNA stability (factors: condition, hours incubated in actinomycin D) and TBK1-ASO reporter specificity (factors: ASO treatment, each reporter) were analyzed with Two-way ANOVA with Dunnett’s post-hoc test where appropriate. Significance is defined as follows: *p < 0.05, **p < 0.01, ***p < 0.001, ****p < 0.0001, and ns, no statistical significance (P ≥ 0.05). All data were analyzed with GraphPad Prism (version 10, USA).

## SUPPLEMENTAL FIGURES

**Figure S1.**
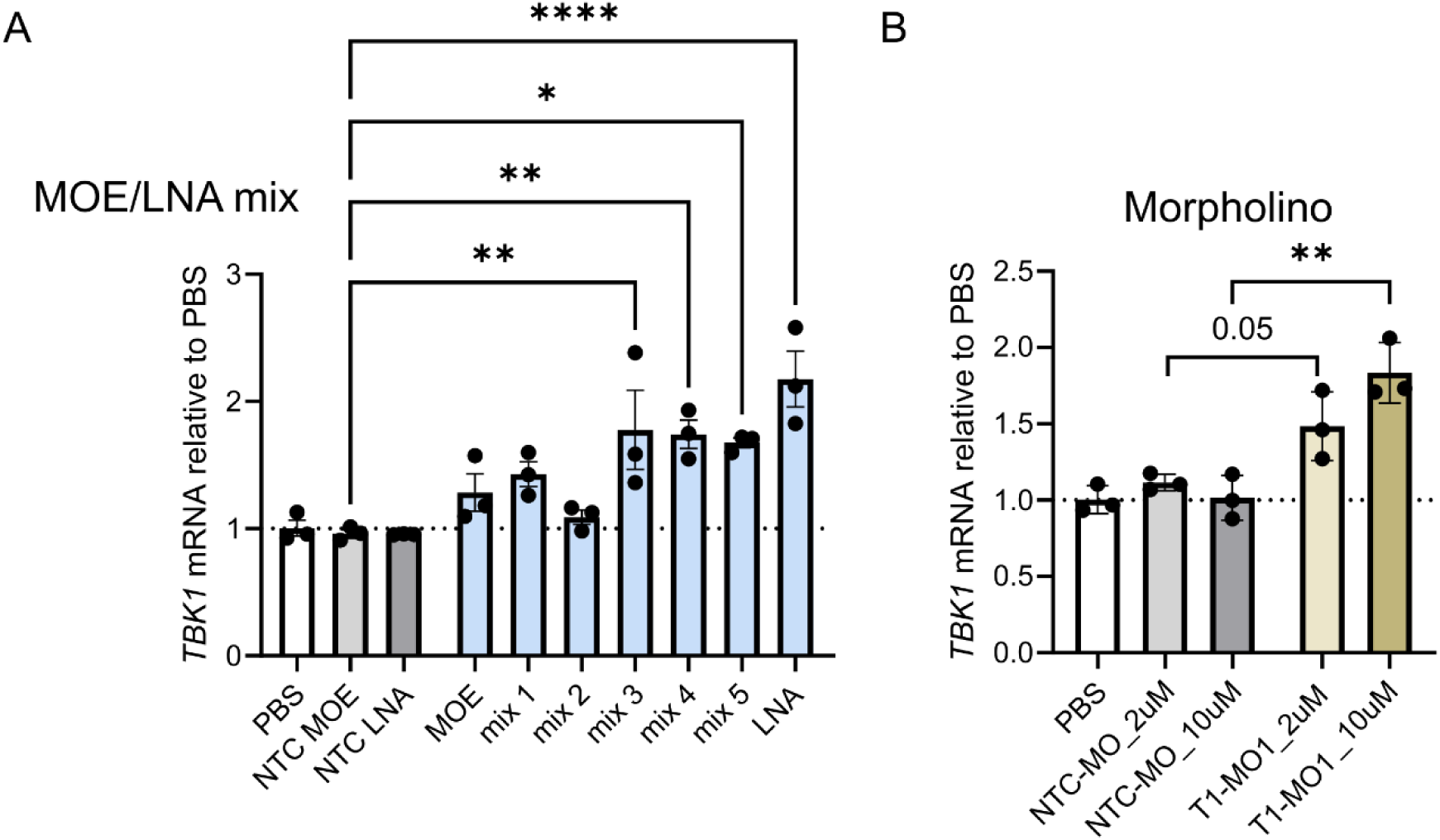
ASO backbone and base chemistries influence TBK1-ASO mRNA upregulation. **(A)** ASOs were developed containing various ratios of methoxyethyl (MOE) or locked nucleic acid (LNA) bases (full MOE, mix 1-5, full LNA). Each was delivered to HEK293 cells alongside a non-targeting control (NTC) MOE or NTC LNA control, and mRNA was quantified by RT-qPCR after 48 hours. Relative abundance of *TBK1* mRNA is shown (ANOVA with Dunnett’s post-hoc, each p<0.05, **p<0.01). **(B)** Morpholinos (MOs) were prepared that matched TBK1-ASO T1 sequence alongside NTC MO control. MOs were delivered to HEK293 cells at 2 uM or 10 uM using Endoporter transfection reagent. *TBK1* mRNA was quantified after 48 hours. (n=3 biological replicates for all experiments) *p<0.05, **p<0.01, ****p<0.0001.

**Figure S2.**
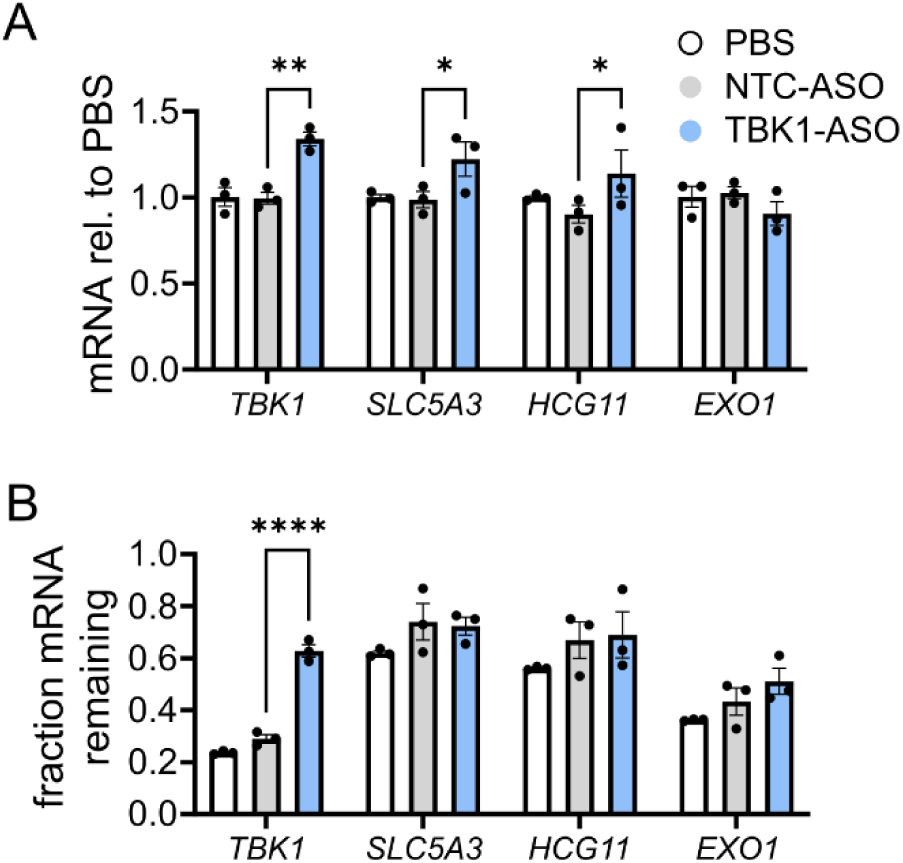
TBK1-ASO does not impact mRNA stability of potential off-target genes. **(A)** The effects of the TBK1-ASO on *TBK1*, *SLC5A3*, *HCG11*, and *EXO1* mRNA were quantified by RT-qPCR (n=3 biological replicates; ANOVA with Dunnett’s post-hoc test, *TBK1*: p=0.002, *SLC5A3*, p=0.03, *HCG11*, p=0.03). **(B)** HEK293 cells were incubated with Actinomycin D for 6 hours, and the fraction of mRNA remaining after 6 hours was quantified by RT-qPCR (n=3 biological replicates; ANOVA with Dunnett’s post-hoc test, *TBK1*: p<0.0001, others: p>0.05). *p<0.05, **p<0.01, ****p<0.0001.

**Figure S3.**
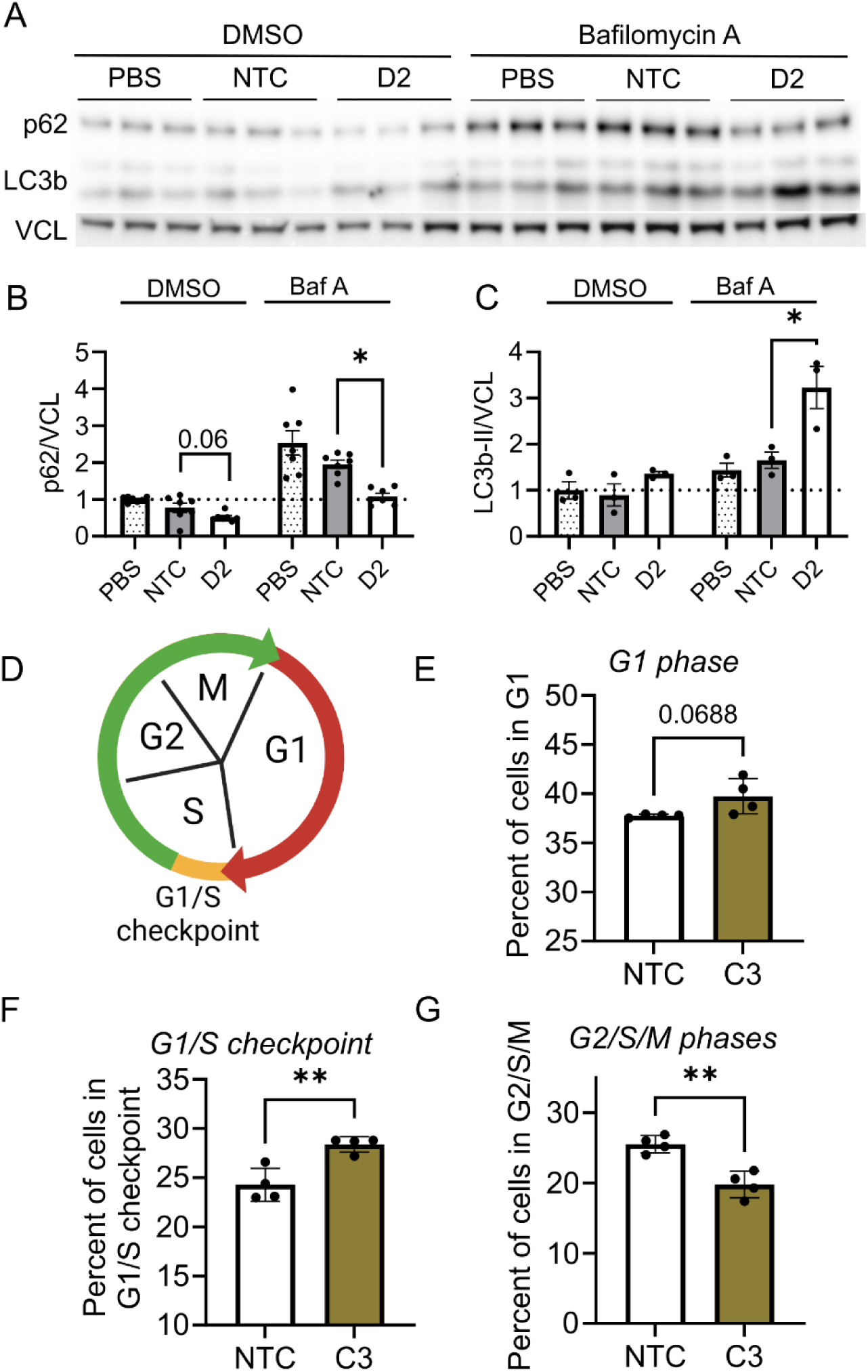
DEPDC5 and CDKN1B upregulating ASOs influence cell states. (A-C) HEK293 cells were treated with 100 nM DEPDC5-ASO D2 for 48 hours, then incubated with Bafilomycin A or DMSO for 6 hours. Protein was collected, and (A) western blot was performed. (B) Relative abundance of p62 normalized to vinculin (VCL) loading control (n=6 biological replicates; ANOVA with Dunnett’s post-hoc test: DMSO: p=0.06, BafA: p=0.01) (C) LC3b-II was normalized to VCL (n=3 biological replicates; ANOVA with Dunnett’s post-hoc test: Baf A: p=0.02). **(D)** The Fucci(SA)5 reporter was stably integrated in HeLa cells and 100 nM NTC or CDKN1B ASO C3 was transfected. After 72 hours, relative fluorescence was measured using flow cytometry, and the proportion of cells in the G1, G1/S transition phase, or the S/G2/M phases was quantified n=4 biological replicates). **(E-G)** The number of cells in the (E) G1 phase showed a trended increase (t-test, p=0.07) alongside a significant increase in cells at the (F) G1/S checkpoint (t-test, p=0.004) and decrease in the (G) G2/S/M phases (t-test, p=0.002). *p<0.05, **p<0.01.

**Figure S4.**
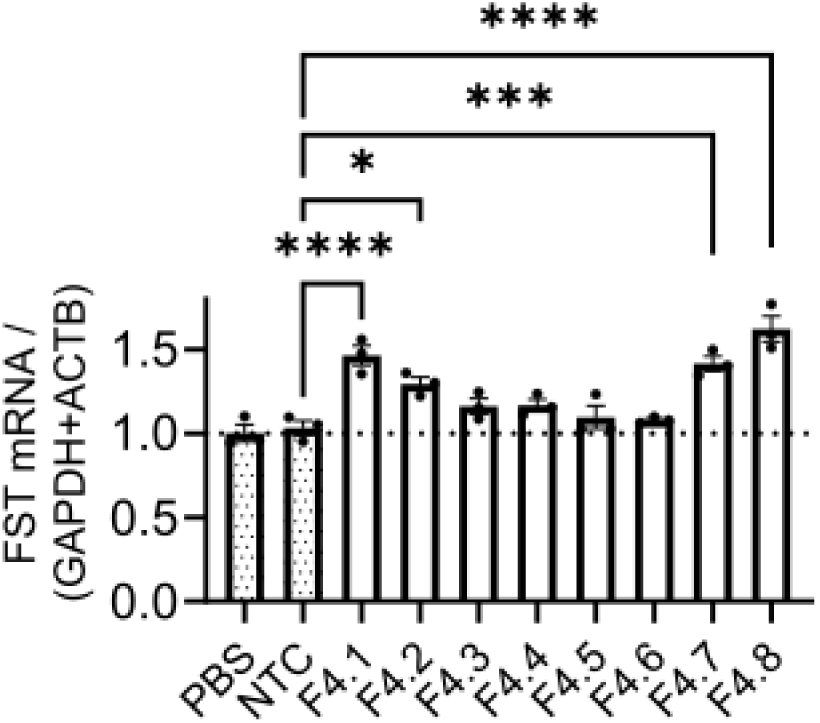
Microwalk of ASOs derived from FST-ASO F4 identifies improved mRNA upregulating candidates. Eight ASOs were developed that targeted near the binding site for FST-ASO F4. These ASOs were delivered to HEK293 cells at 100 nM, and after 48 hours, RNA was collected for RT-qPCR analysis for *FST* mRNA. (n=3 biological replicates per condition; ANOVA with Dunnett’s post-hoc: p<0.05). *p<0.05, ***p<0.001, ****p<0.0001.

## ACKNOWLEDGMENTS

The authors thank the Genome Technology Access Center at the McDonnell Genome Institute (GTAC@MGI) and the High Throughput Computing Facility at the Center for Genome Sciences and Systems Biology. The authors thank Dr. Simona Sarafinovska for critical reading.

## AUTHOR CONTRIBUTIONS

Conceptualization: BDB, KMS, SD, TMM

Methodology: BDB, AN, MH, DAG, CFJ

Investigation: BDB, AN, CFF

Visualization: BDB, AN

Writing: BDB and TMM

Project coordination: BDB, JDD, KMS, SD, TMM

## REFERENCES

1. Huang, N., Lee, I., Marcotte, E. M. & Hurles, M. E. Characterising and Predicting Haploinsufficiency in the Human Genome. PLOS Genet. 6, e1001154 (2010).

2. Freischmidt, A. et al. Haploinsufficiency of TBK1 causes familial ALS and fronto-temporal dementia. Nat. Neurosci. 18, 631–636 (2015).

3. Mahboob, M., Rout, P., Leslie, S. W. & Bokhari, S. R. A. Autosomal Dominant Polycystic Kidney Disease. in StatPearls (StatPearls Publishing, Treasure Island (FL), 2025).

4. Anisimova, A. S., Alexandrov, A. I., Makarova, N. E., Gladyshev, V. N. & Dmitriev, S. E. Protein synthesis and quality control in aging. Aging 10, 4269–4288 (2018).

5. Guo, J. et al. Aging and aging-related diseases: from molecular mechanisms to interventions and treatments. Signal Transduct. Target. Ther. 7, 1–40 (2022).

6. Corey, D. R., Damha, M. J. & Manoharan, M. Challenges and Opportunities for Nucleic Acid Therapeutics. Nucleic Acid Ther. 32, 8–13 (2022).

7. Sun, X., Setrerrahmane, S., Li, C., Hu, J. & Xu, H. Nucleic acid drugs: recent progress and future perspectives. Signal Transduct. Target. Ther. 9, 1–31 (2024).

8. Becker, K. et al. Quantifying post-transcriptional regulation in the development of Drosophila melanogaster. Nat. Commun. 9, 4970 (2018).

9. Gebauer, F. & Hentze, M. W. Molecular mechanisms of translational control. Nat. Rev. Mol. Cell Biol. 5, 827–835 (2004).

10. Bohn, J. A. et al. Identification of diverse target RNAs that are functionally regulated by human Pumilio proteins. Nucleic Acids Res. 46, 362–386 (2018).

11. Boros, B. D., Schoch, K. M., Kreple, C. J. & Miller, T. M. Antisense Oligonucleotides for the Study and Treatment of ALS. Neurother. J. Am. Soc. Exp. Neurother. 19, 1145–1158 (2022).

12. Collotta, D., Bertocchi, I., Chiapello, E. & Collino, M. Antisense oligonucleotides: a novel Frontier in pharmacological strategy. Front. Pharmacol. 14, 1304342 (2023).

13. Bennett, C. F., Baker, B. F., Pham, N., Swayze, E. & Geary, R. S. Pharmacology of Antisense Drugs. Annu. Rev. Pharmacol. Toxicol. 57, 81–105 (2017).

14. Davis, S., Lollo, B., Freier, S. & Esau, C. Improved targeting of miRNA with antisense oligonucleotides. Nucleic Acids Res. 34, 2294–2304 (2006).

15. Davis, S. et al. Potent inhibition of microRNA in vivo without degradation. Nucleic Acids Res. 37, 70–77 (2009).

16. Hedaya, O. M. et al. Secondary structures that regulate mRNA translation provide insights for ASO-mediated modulation of cardiac hypertrophy. Nat. Commun. 14, 6166 (2023).

17. Li, Y. et al. Targeting 3′ and 5′ untranslated regions with antisense oligonucleotides to stabilize frataxin mRNA and increase protein expression. Nucleic Acids Res. 49, 11560– 11574 (2021).

18. Liang, X. et al. Translation efficiency of mRNAs is increased by antisense oligonucleotides targeting upstream open reading frames. Nat. Biotechnol. 34, 875– 880 (2016).

19. Lim, K. H. et al. Antisense oligonucleotide modulation of non-productive alternative splicing upregulates gene expression. Nat. Commun. 11, 3501 (2020).

20. Kota, J. et al. Follistatin Gene Delivery Enhances Muscle Growth and Strength in Nonhuman Primates. Sci. Transl. Med. 1, 6ra15 (2009).

21. Brockmann, K. et al. Autosomal dominant Glut-1 deficiency syndrome and familial epilepsy. Ann. Neurol. 50, 476–485 (2001).

22. Ishida, S. et al. Mutations of DEPDC5 cause autosomal dominant focal epilepsies. Nat. Genet. 45, 552–555 (2013).

23. Ison, H. E., Clarke, S. L. & Knowles, J. W. Familial Hypercholesterolemia. in GeneReviews® (eds. Adam, M. P. et al.) (University of Washington, Seattle, Seattle (WA), 1993).

24. Lee, S.-J. et al. Regulation of Muscle Mass by Follistatin and Activins. Mol. Endocrinol. 24, 1998–2008 (2010).

25. Li, R. et al. ITPR1 variant-induced autosomal dominant hereditary spastic paraplegia in a Chinese family. Front. Neurol. 15, (2024).

26. Yamada, T. et al. Systematic Analysis of Targets of Pumilio-Mediated mRNA Decay Reveals that PUM1 Repression by DNA Damage Activates Translesion Synthesis. Cell Rep. 31, 107542 (2020).

27. Cottrell, K. A., Chaudhari, H. G., Cohen, B. A. & Djuranovic, S. PTRE-seq reveals mechanism and interactions of RNA binding proteins and miRNAs. Nat. Commun. 9, 301 (2018).

28. Watts, J. K. & Corey, D. R. Silencing disease genes in the laboratory and the clinic. J. Pathol. 226, 365–379 (2012).

29. Roberts, T. C., Langer, R. & Wood, M. J. A. Advances in oligonucleotide drug delivery. Nat. Rev. Drug Discov. 19, 673–694 (2020).

30. Schoch, K. M. & Miller, T. M. Antisense Oligonucleotides: Translation from Mouse Models to Human Neurodegenerative Diseases. Neuron 94, 1056–1070 (2017).

31. Roberts, T. C., Langer, R. & Wood, M. J. A. Advances in oligonucleotide drug delivery. Nat. Rev. Drug Discov. 19, 673–694 (2020).

32. Feichtenschlager, V. et al. Suppression of NRAS-mutant melanoma growth with NRAS-targeting Antisense Oligonucleotide treatment reveals therapeutically relevant kinase co-dependencies. Commun. Med. 5, 216 (2025).

33. Magner, D., Biala, E., Lisowiec-Wachnicka, J. & Kierzek, R. Influence of mismatched and bulged nucleotides on SNP-preferential RNase H cleavage of RNA-antisense gapmer heteroduplexes. Sci. Rep. 7, 12532 (2017).

34. Dibbens, L. M. et al. Mutations in DEPDC5 cause familial focal epilepsy with variable foci. Nat. Genet. 45, 546–551 (2013).

35. Bencivenga, D. et al. A cancer-associated CDKN1B mutation induces p27 phosphorylation on a novel residue: a new mechanism for tumor suppressor loss-of-function. Mol. Oncol. 15, 915–941 (2021).

36. Le Toriellec, E. et al. Haploinsufficiency of CDKN1B contributes to leukemogenesis in T-cell prolymphocytic leukemia. Blood 111, 2321–2328 (2008).

37. Ando, R., Sakaue-Sawano, A., Shoda, K. & Miyawaki, A. Two coral fluorescent proteins of distinct colors for sharp visualization of cell-cycle progression. Cell Struct. Funct. 48, 135–144 (2023).

38. Tang, R. et al. Gene therapy for follistatin mitigates systemic metabolic inflammation and post-traumatic arthritis in high-fat diet-induced obesity. Sci. Adv. 6, eaaz7492 (2020).

39. Winbanks, C. E. et al. Follistatin-mediated skeletal muscle hypertrophy is regulated by Smad3 and mTOR independently of myostatin. J. Cell Biol. 197, 997–1008 (2012).

40. Overby, S. J. et al. Proof of concept of peptide-linked blockmiR-induced MBNL functional rescue in myotonic dystrophy type 1 mouse model. Mol. Ther. Nucleic Acids 27, 1146–1155 (2022).

41. Ting, K. K. et al. Therapeutic regulation of VE-cadherin with a novel oligonucleotide drug for diabetic eye complications using retinopathy mouse models. Diabetologia 62, 322– 334 (2019).

42. Dao, K., Jungers, C. F., Djuranovic, S. & Mustoe, A. M. U-rich elements drive pervasive cryptic splicing in 3’ UTR massively parallel reporter assays. BioRxiv Prepr. Serv. Biol. 2024.08.05.606557 (2024) doi:10.1101/2024.08.05.606557.

43. Kedersha, N. L., Gupta, M., Li, W., Miller, I. & Anderson, P. RNA-binding proteins TIA-1 and TIAR link the phosphorylation of eIF-2 alpha to the assembly of mammalian stress granules. J. Cell Biol. 147, 1431–1442 (1999).

44. Lin, W.-J., Duffy, A. & Chen, C.-Y. Localization of AU-rich element-containing mRNA in cytoplasmic granules containing exosome subunits. J. Biol. Chem. 282, 19958–19968 (2007).

45. Sela-Brown, A., Silver, J., Brewer, G. & Naveh-Many, T. Identification of AUF1 as a Parathyroid Hormone mRNA 3′-Untranslated Region-binding Protein That Determines Parathyroid Hormone mRNA Stability*. J. Biol. Chem. 275, 7424–7429 (2000).

46. Kedde, M. et al. A Pumilio-induced RNA structure switch in p27-3′ UTR controls miR-221 and miR-222 accessibility. Nat. Cell Biol. 12, 1014–1020 (2010).

47. Zhou, Y., Ng, D. Y., Richards, A. M. & Wang, P. Loss of full-length pumilio 1 abrogates miRNA-221-induced gene p27 silencing-mediated cell proliferation in the heart. Mol. Ther. Nucleic Acids 27, 456–470 (2021).

48. Phillips, D. J. & de Kretser, D. M. Follistatin: a multifunctional regulatory protein. Front. Neuroendocrinol. 19, 287–322 (1998).

49. Mendell, J. R. et al. A Phase 1/2a Follistatin Gene Therapy Trial for Becker Muscular Dystrophy. Mol. Ther. 23, 192–201 (2015).

50. Ge, G. et al. Long-term benefits of hematopoietic stem cell-based macrophage/microglia delivery of GDNF to the CNS in a mouse model of Parkinson’s disease. Gene Ther. 31, 324–334 (2024).

51. Grunewald, M. et al. Counteracting age-related VEGF signaling insufficiency promotes healthy aging and extends life span. Science 373, eabc8479 (2021).

52. Ando, R., Sakaue-Sawano, A., Shoda, K. & Miyawaki, A. Two coral fluorescent proteins of distinct colors for sharp visualization of cell-cycle progression. Cell Struct. Funct. 48, 135–144 (2023).

